# Methylation of 23S rRNA G748 and the ribosomal protein L22 Lys-94 are critical factors for maintaining the association between ribosome stalling and proteome composition in *Streptococcus pneumoniae*

**DOI:** 10.1101/2021.04.14.439922

**Authors:** Tatsuma Shoji

**Author notes:** Correspondence should be addressed to T.S.

## Abstract

**Background:** 23S rRNA modification located at the nascent peptides exit tunnel plays an important role in both translation processes and the binding of the antibiotics. Methylation of the guanine at position 748 (m^1^G748) in 23S rRNA in *Streptococcus pneumoniae* is involved in the ribosome stalling and the binding of the antibiotic telithromycin (TEL). The disruption of the gene encoding RlmA^II^ which methylates 23S rRNA G748 results in the increased resistance of TEL in *S. pneumoniae*. However, an isolated high-level TEL-resistant *S. pneumoniae* strain indicated that additional undescribed factors were involved in TEL resistance in *S. pneumoniae*.

**Results:** We successfully isolated a high-level TEL-resistant *S. pneumoniae* RlmA^II^ mutant and determined the whole-genome sequence. The lysine residue at the position 94 in ribosomal protein 22 (L22 K94) was critical in binding of TEL to the ribosome. A growth competition assay showed that L22 K94 was required for the function of m^1^G748. Ribosome profiling revealed that m^1^G748 and L22 K94 were both essential to maintain the relationship between the ribosome stalling and proteome composition.

**Conclusion:** In *S. pneumoniae*, the combination of methylation status of G748 and the residue at position 94 in L22 are essential for both the distribution of ribosome stalling and the binding of TEL to ribosomes.

## 1. Background

Ribosomes are the cellular location where proteins are synthesized. The structure of the peptidyltransferase center (PTC) is essential for the protein translation reaction. This structure consists of not only 23S rRNA but also the result of the biochemical interaction between the nascent peptides and the nascent peptides exit tunnel (NPET) [1,2]. Therefore, the structure of the NPET is believed to include ribosomal protein and rRNA, and rRNA modification is critical for the function of the ribosome in translation of mRNA to protein [3]. Previously, we demonstrated that ribosome stalling is affected by the methylation of the 23S rRNA at position G748, which is located at the NPET near the PTC [4].

Several modifications located in the NPET play important roles in determining antibiotic resistance or sensitivity [5]. Previously, we showed that the inactivation of the methyltransferase RlmA^II^, which methylates the N-1 position of nucleotide G748 (m^1^G748), or the inactivation of the methyltransferase RlmCD, which methylates the N-5 position of nucleotide U747 (m^5^U747) and C-5 position of U1939, results in increased resistance to telithromycin (TEL) in *erm*(B)-carrying *Streptococcus pneumonia* [6,7]. The minimum inhibitory concentration (MIC) for TEL in the RlmA^II^-disrupted or RlmCD-disrupted *S. pneumoniae* mutants is 16 to 32 or 8 μg/mL, respectively. However, Walsh et al. isolated a high-level TEL-resistant *S. pneumoniae* whose MIC for TEL was more than 512 μg/mL, [8] indicating that additional factors associated to the TEL-resistance are present other than m^1^G748 and m^5^U747.

In this study, we sought to identify the additionally factors that are associated with the high-level TEL resistance in *S. pneumoniae*. We initially isolated a high-level TEL-resistant *S. pneumoniae* mutant from an RlmA^II^ mutant and determined the whole-genome sequence of these mutants. Subsequent analysis showed that the lysine residue at position 94 in the ribosomal protein L22 (L22 K94) is a key factor involved in high-level TEL resistance and that L22 K94 is necessary for the function of m^1^G748.

## 2. Results

### 2.1. Mutation of L22 K94E produces high-level TEL resistance in *S. pneumoniae*

We previously isolated TEL-resistant mutants from S1, a clinically isolated *S. pneumoniae* strain with low TEL susceptibility (MIC, 2 μg/mL), by selection with 8 and 16 μg/mL of TEL to yield 10 and 6 mutants, respectively. Each of the 16 strains (Sp32 to Sp47) contained one unique nucleotide difference in the *tlrB* gene that encodes RlmA^II^ but did not contain any shared mutations [6]. This result indicates that mutations involving high-level TEL resistance would occur only after the mutation of the *tlrB* gene. Thus, we sought to isolate high-level TEL resistance mutants from Sp36, one of 16 mutants described above, by selection with 128 μg/mL of TEL. As a result, several mutants (Sp48 to Sp52) were isolated that all increased MICs (> 512 μg/mL).

To determine the mutations involved in high-level TEL resistance, whole-genome sequencing of strains S1 and Sp52 was performed by next-generation sequencing technology, and open reading frame (ORF) sequences were analyzed for the two strains. As a result, one mutation was found in a gene encoding a ribosomal protein L22, leading to an Lys94Glu alteration in this protein in Sp52. Furthermore, this mutation was confirmed in mutants Sp48 to Sp51 but was absent in mutants Sp32 to Sp47. According to the three-dimensional crystal structure of the *Escherichia coli* 50S subunit of the 70S ribosome bound to TEL (Protein Data Bank: 3OAT) [9], the L22 K90 mutant protein, which corresponds to L22 K94 in *S. pneumoniae,* is a component of NPET and is located opposite m^1^G748; this residue appears to form three hydrogen bonds to the oxygen atom in TEL via the methyl group at the end of side chain in K90 (Figure 1). Therefore, the K94E mutation would lose these hydrogen bonds. These results suggest that the L22 K94E mutation leads to the high level of TEL resistance in *S. pneumoniae*.

**Fig. 1.**
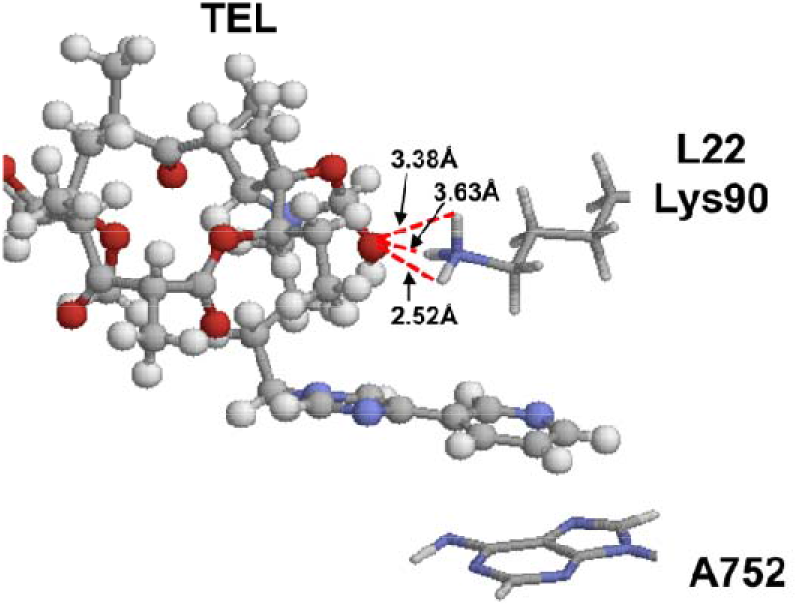
Structure of TEL binding to the ribosome 50S subunit. The TEL molecule is shown by a stick-and-ball model. The residues of 23S rRNA A752 and L22 K94 are shown. Dashed lines show the distances from the oxygen atom in TEL to the hydrogen atom of the amino group in the side chain of L22 K94.

### 2.2. L22 K94 is necessary for the function of m^1^G748

We were unable to isolate L22 K94E mutant directly from S1, which implies a genetic interaction between m^1^G748 and L22 K94. The *S. pneumonia* mutant that harbors the wild-type *tlrB* and the mutated L22 K94E may exhibit a lower fitness compared with that of S1, Sp36, and Sp52. To examine this hypothesis, we introduced a plasmid encoding the wild-type *tlrB* gene to Sp52 (Sp375) and compared the growth for the following four strains: S1, Sp36, Sp52, and Sp375 (Figure 2A). As expected, Sp52 and Sp375 exhibited much lower growth compared with that of S1 and Sp36. To clarify the growth difference between Sp52 and Sp375, we performed a growth competition assay between these two strains. As a result, Sp375 rapidly disappeared during subculturing (Figure 2B). This result clearly demonstrated that L22 K94 is required for the functioning of m^1^G748.

**Fig. 2.**
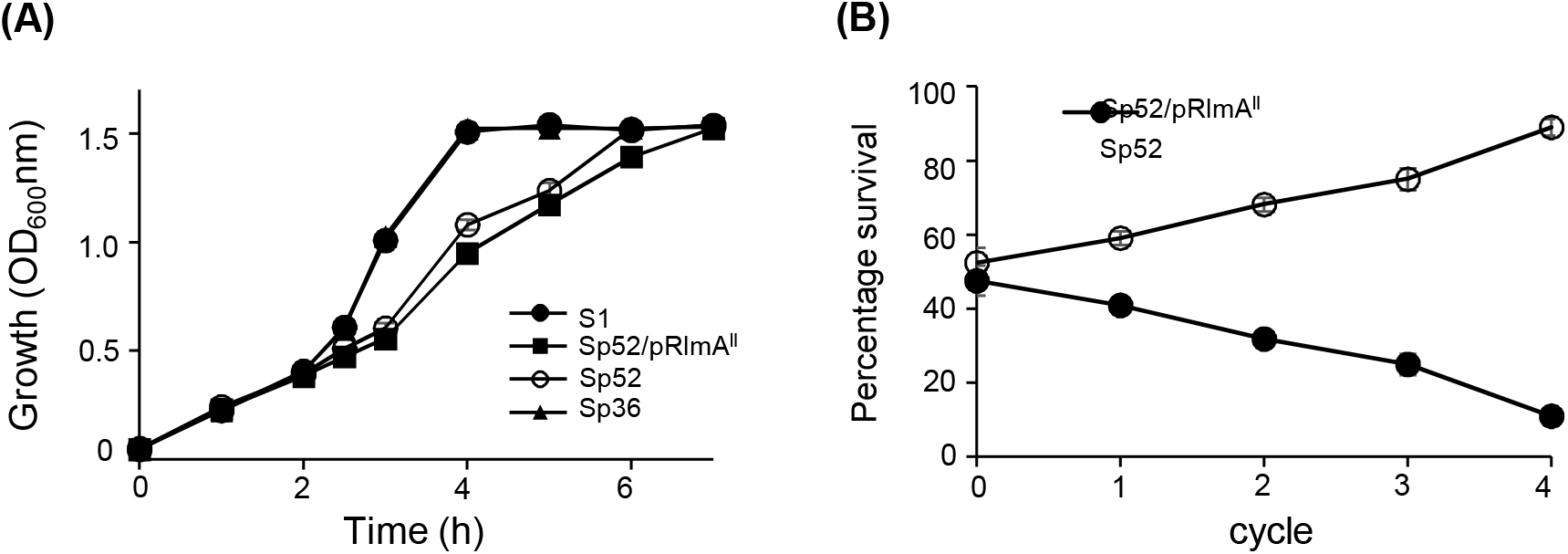
Effect of m^1^G748/L22 K94 on growth. (A) Growth of strains S1 (closed circle), Sp375 (Sp52/pRlmA^II^; closed square), Sp52 (open square), and Sp36 (closed triangle) in BHI-Y broth at 37°C and 5% CO_2_. (B) Growth competition between Sp375 (Sp52/pRlmA^II^; open circle) and Sp52 (closed circle). Sp375 was mixed with an equal quantity of Sp52 cells and grown as described in Materials and Methods. The Sp375 cells were assayed by plating on BHI-Y agar supplemented with 5% horse blood plus kanamycin, whereas the sum of the Sp375 and Sp52 cells were obtained by plating on BHI-Y agar supplemented with 5% horse blood. The percent of Sp375 or Sp52 cells remaining at each sub-culture was calculated, and the percent abundance of each strain was plotted. Dilution between cycles was 1:20. Mean values and standard deviations from at least three independent experiments are given in (A) and (B).

### 2.3. m^1^G748/L22 K94 affects ribosome stalling

We previously showed that m^1^G748 affects ribosome stalling. Thus, to investigate the role of L22 K94 in ribosome stalling, we performed ribosome profiling assay in four strains: WT, S1ΔrlmA^II^/pRlmA^II^(wild-type); L22 mutant, Sp52ΔrlmA^II^/pRlmA^II^(wild-type); Double mutant, Sp52ΔrlmA^II^/pRlmA^II^(C23R); RlmA^II^ mutant, S1ΔrlmA^II^/pRlmA^II^(C23R); note that RlmA^II^ with a mutation of C23R exhibits no methyltransferase activity [6]. The *erm*(B) operon, where the ribosomes stall at the *ermBL* region [10], was selected as an example to observe the distribution of ribosomes. The ribosome position and density in the *erm*(B) operon were completely different between the four strains (Figure 3A).

**Fig. 3.**
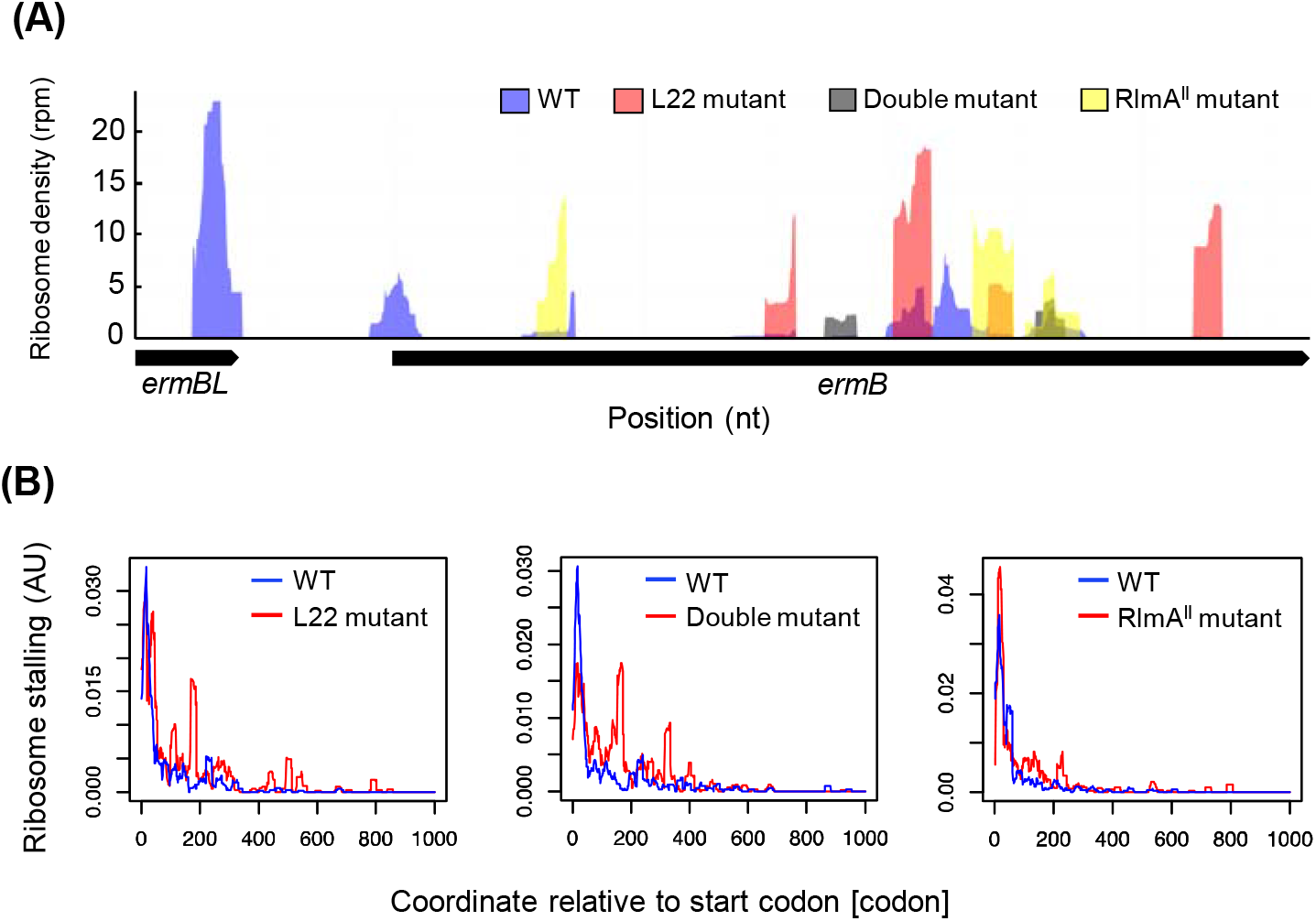
Ribosome profiling in *S. pneumoniae* reveals ribosome stalling. (A) Ribosome density plot of the *erm*(B) operon shows the difference in ribosome occupancy among WT, L22 mutant, Double mutant, and RlmA^II^ mutant strains. (B) Metagene profile of the ribosome stalling in WT or L22 mutant (left panel), WT or Double mutant (middle panel), and WT or RlmA^II^ mutant (right panel). The A-site density across the stalling genes was aligned relative to the start codon.

The difference in the ribosome occupancy among these four strains in the *erm*(B) operon led us to speculate on the general role of m^1^G748 and L22 K94 in ribosome stalling. We previously defined ribosome stalling as a A-site density (see Method section) higher than eight based on the result of the RNA footprinting assay [4]. Thus, we performed metagene analysis to obtain an overview of global differences in ribosome stalling across all ORFs (Figure 3B). This showed that ribosome stalling across all ORFs in mutants was higher than that of WT especially around the 100-500 residue region, indicating the important role of m^1^G748 and L22 K94 in ribosome stalling.

### 2.4. m^1^G748/L22 K94 has a major impact on the interaction between the NPET and nascent peptides

One of the reasons for ribosome stalling is the biochemical interaction between nascent peptides and the NPET. To investigate the role of m^1^G748 and L22 K94 on the biochemical interaction in NPET, the “stalling peptides” were identified as previously described [11]. Briefly, we first collected the nascent peptide sequences in the exit tunnel for all of ribosome stallings and calculated the probability of the occurrence for each of 8,000 tripeptides. Then, we defined the stalling peptides as tripeptides with a probability higher than 0.9999 and compared the stalling peptides among the four strains. Surprisingly, there were very few stalling peptides that were commonly found among these strains (Figure 4A), suggesting that the biochemical interaction in the NPET differs among the four strains.

**Fig. 4.**
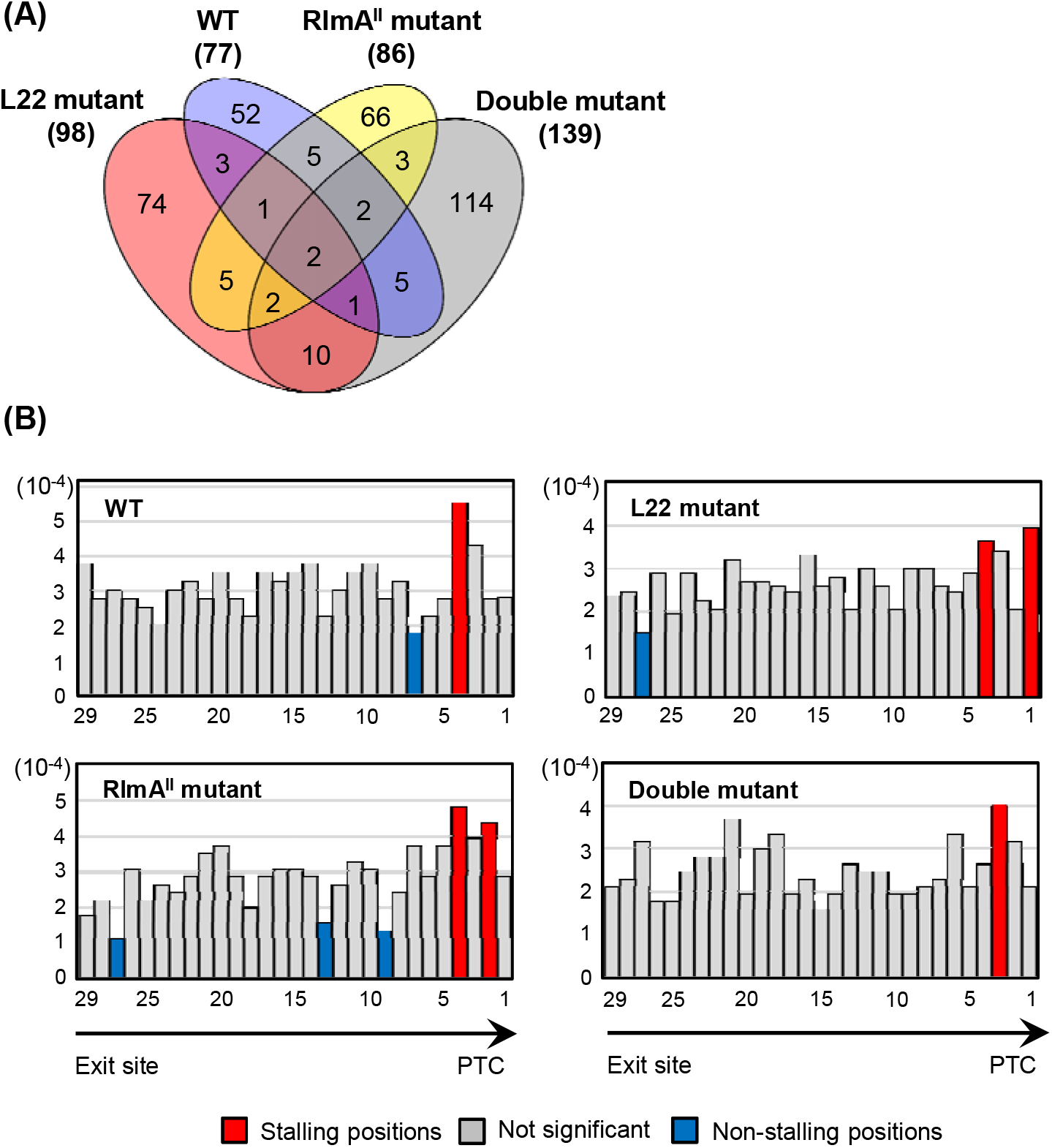
Effect of m^1^G748/L22 K94 on the interactions between the ribosomal exit tunnel and the nascent peptide. (A) Venn diagram for stalling peptide sequences in WT, L22 mutant, Double mutant, and RlmA^II^ mutant strains. The number of stalling peptide sequences are shown in parentheses. (B) The average distribution of stalling peptide sequences along the length of the tunnel. The positions along the tunnel (x-axis) represent the first position the peptide occupies. The arrow underneath the x-axis stands for the direction of NPET. The height of the bar is the probability that this position is occupied (average probability over all peptide sequences defined in the title). Bars are colored red/blue if their corresponding probabilities were significantly (P < 0.05) higher/lower than random, respectively; other bars (P > 0.05) appear in gray.

The length of the NPET is approximately 31 amino acids [11]. To test whether m^1^G748 and L22 K94 directly or indirectly interact with the NPET, the frequency of the occurrence of the stalling peptides at each of the 29 positions in the NPET was calculated. The stalling peptides tend to occur near the PTC in all four strains (Figure 4B), indicating a direct biochemical interaction between m^1^G748/L22 K94 and NPET.

### 2.5. m^1^G748/L22 K94 is a general factor which affects the association between ribosome stalling and proteome composition

Stalling peptides cause ribosome stalling, which essentially ends the translation process [12, 13]. Therefore, the sequences of the stalling peptides are expected to be evolutionary underrepresented. To examine this hypothesis, we first identified 360 overrepresented and 382 underrepresented tripeptides in the proteome of *S. pneumoniae*. Then, the enrichment of over- and underrepresented tripeptides in the stalling peptides was calculated using the Fisher's exact test (Figure 5A). While both over- and underrepresented peptides were significantly enriched in the stalling peptides set of WT, neither of these sets were enriched in the other three strains. This indicated that the stalling peptides are evolutionarily conserved and that m^1^G748 and L22 K94 are important for maintaining the relationship between ribosome stalling and the proteome composition.

**Fig. 5.**
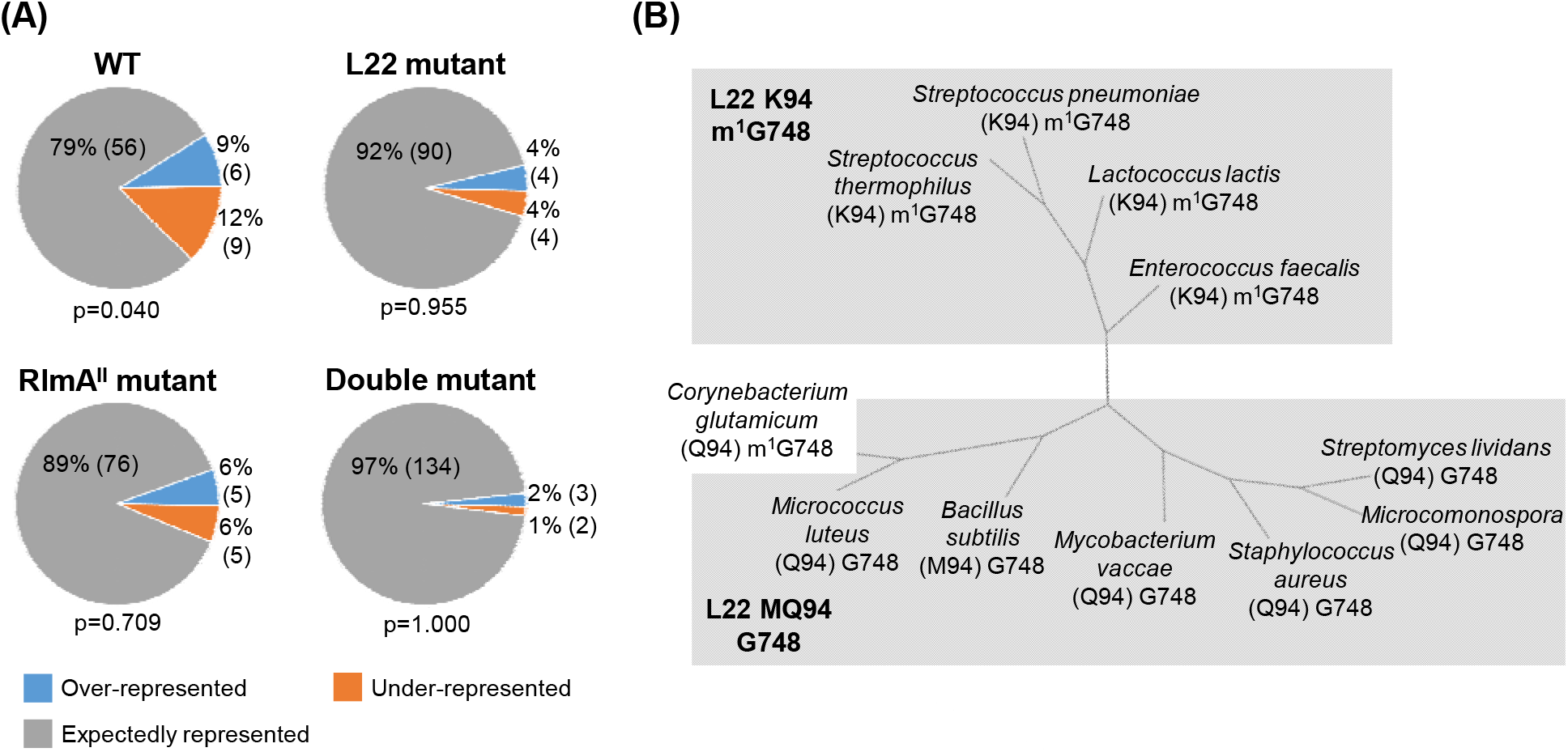
Role of m^1^G748/L22 K94 in the association between ribosome stalling and proteome composition. (A) Distribution of the number of stalling peptide sequences with respect to proteome representation in WT, L22 mutant, Double mutant, and RlmA^II^ mutant; the P-value represents the statistical significance of the over- and underrepresented proportion. (B) A phylogenetic tree based on average linkage clustering of the selected gram-positive bacteria using the random vector produced by proteome representation. The proteome representation segregated into two groups in a manner largely consistent with the combination of the residue at position 90 in L22 and the status of G748 methylation. The amino acids residues at position 90 in L22 are shown in parentheses after the name of the organism followed by the status of G748 methylation.

To further examine the role of m^1^G748 and L22 K94 in the proteome composition, we calculated the 8,000-dimension vector representing the frequency of the occurrence of all 8,000 tripeptides in the proteome for 12 gram-positive bacterial species in which the methylation of G748 and the sequence of L22 have been revealed [14]. Then, we drew a phylogenetic tree based on this frequency vector. Surprisingly, the proteome composition segregated into two groups in a manner largely consistent with the combination of the mutated residue at position 94 in L22 and the status of G748 methylation (Figure 5B), suggesting that m^1^G748/L22 K94 are the general factors for controlling the ribosome stalling against the proteome composition.

## 3. Discussion

In this study, we successfully identified L22 K94 as the key factor involved in the high-level TEL resistance in *S. pneumoniae*. We revealed that L22 K94 is required for the function of m^1^G748, which is important for ribosome stalling at the appropriate position to maintain proteome composition. The reason TEL exhibits a strong effect specifically on *S. pneumoniae*, and why TEL-resistant mutants of *S. pneumoniae* are rarely clinically isolated [15] may be because the regions where TEL binds are indispensable for *S. pneumoniae* to control ribosome stalling at the correct position. Here, the growth of both the Double mutant and the L22 mutant were much lower than that of the others (Figure 2).

L22 K94 is located on the opposite side of m^1^G748 in the NPET [9]. Several rRNA modifications are clustered around m^1^G748 and L22 K94 [16]. These modifications, together with m^1^G748 and L22 K94, compose a narrow gate in the NPET called the “discriminating gate” [1]. Thus, the discriminating gate may function to control ribosome stalling at the appropriate position to maintain proteome composition. This implies the structure of the discriminating gate might be different among species since the discriminating gate is responsible for the proteome composition which is different among species. Therefore, antibiotics that target the discriminating gate would be species-specific and would not induce resistance in non-targeted bacteria. This is a new concept for designing antibiotics, and therefore, discriminating gate-targeting antibiotics may form a new class of antibiotics that should be investigated in future studies.

The combination of the residue at the position 94 in L22 and the status of G748 methylation correlated to the composition of the proteome (Figure 5B). This means the distribution of the ribosome stalling could explain the differences among species. Since the distribution of the ribosome stalling is summarized as the probability of the occurrence for each 8,000 tripeptide, or more generally each 20^n^ n-length peptide, each species could be represented mathematically by using the 20^n^ dimension vector. We suggest that this 20^n^ vector “stalling vector” would be very useful as a mathematical representation of the species. For example, the distance between two species is easily calculated based on the definition of the distance in mathematics without defining the distance between nucleotides, in which the rational for the definition is unclear. Furthermore, since the distribution of the ribosome stalling depends on the cell condition or experiment condition [10], the stalling vector could be used as the biological definition for the cell or experiment condition.

## 4. Conclusions

We identified that L22 K94 was a key factor involved in the high-level TEL resistance in *S. pneumoniae* and demonstrated that the residue at position 94 in L22 and the status of the methylation of G748 are essential to maintain the appropriate ribosome stalling for the proteome composition. Therefore, this region in the NPET may be a new target for a novel class of antibiotics.

## 5. Methods

### 5.1. Bacterial strains, plasmids, and media

Bacterial strains and plasmids are shown in Supplementary Table S1 and S2, respectively. *S. pneumoniae* strain S1 with reduced TEL susceptibility (MIC, 2 μg/mL) was clinically isolated in Japan [6]. Pneumococci were routinely cultured at 37°C and 5% CO_2_ in brain-heart infusion media with 0.5% yeast extract (BHI-Y) broth and BHI-Y agar, supplemented with 5% horse blood. *E. coli* was grown in L broth (1% Bact-tryptone, 0.5% Bact yeast extract, and 0.5% sodium chloride, pH 7.4) and L agar. When necessary, the medium was supplemented with kanamycin (25–600 μg/mL), spectinomycin (100 μg/mL), and ampicillin (25 μg/mL).

### 5.2. Transformation

Synthetic competence-stimulating peptide (CSP) 1 and the method of Iannelli and Pozzi [17] were used to produced transformation-competent *S. pneumoniae* S1.

### 5.3. Antimicrobial susceptibility testing

Susceptibility to antibiotics was determined by the serial two-fold dilution method, using Mueller-Hinton agar plates supplemented with 5% lysed horse blood. Susceptibility or resistance of pneumococci to TEL was assessed in accordance with the recommendations of the Clinical and Laboratory Standards Institute [18].

### 5.4. Next-generation sequencing and data analysis

Relative mutations in the bacterial genome were estimated by sequencing the genomic DNA samples extracted from each strain. Total DNA were extracted using the method of Blue and Mitchell [19]. For preparation of library DNA, 500 ng of total DNA was sheared to 800 bp by Covaris (M&S). The sheared DNA products were purified using a MiniElute PCR purification kit (Qiagen) and were then used as templates for pyrosequencing with a GS Junior platform (Roche). The data were treated by a data analysis pipeline. Reads from both S1 and Sp52 were assembled into contigs by a de novo assembler. The nucleotide sequence polymorphisms among these strains were listed by mapping sequence reads on the assembled contigs with a reference mapper program. We collected nucleotide substitutions meeting the following two conditions: (i) the substitution was supported by five sequencing reads, and (ii) 90% of the reads supported the substitution.

### 5.5. RNA-Seq

*S. pneumoniae* cultures were grown to log-phase, and 2.8 mL of cultures were then added to 2.8 mL of RNA Lysis Buffer (1% SDS, 0.1 M NaCl, and 8 mM EDTA) pre-heated to 100°C and vortexed for 2 min. The resulting lysates were added to the 5.6 mL of 100°C-preheated acid phenol (Sigma-Aldrich) and vortexed for 5 min. After centrifuging, RNA was extracted from the aqueous phase using DirectZol (Zymo Research). rRNA was isolated from the total RNA using MICROBExpress (Ambion). The resulting total mRNA (400 ng as an input) was used for constructing the DNA library using a KAPA Stranded RNA-Seq Library Preparation Kit Illumina platforms (KK8400). DNA libraries were sequenced using the Illumina HiSeq 1500 system with single-end reads. The Illumina libraries were preprocessed by clipping the Illumina adapter sequence using Trimmomatic v.0.39 [20] and then aligned to the S1 genome sequence^6^ using HASAT2 v.2.2.1 [21].

### 5.6. Ribo-Seq

Libraries were prepared as previously described in [4]. *S. pneumoniae* cultures were grown to log-phase. Next, cells were pretreated for 2 min with 100 μg/mL chloramphenicol before being pelleted by centrifugation. After decanting the supernatant, the cell pellets were resuspended in 2.5 mL of resuspension buffer (10 mM MgCl_2_, 100 mM NH_4_Cl, 20 mM Tris, pH 8.0, and 1 mM chloramphenicol). Then, cells were sonicated on ice and centrifuged, and 25 Abs_260_ ribosome units (1 A_260_ = 12 μg/μL) were digested with MNase (Roche). Digested samples were carefully loaded onto sucrose gradients and centrifuged at 124,700 × *g* for 8 hr at 4°C. After centrifugation, ribosome footprint-associated fractions were pooled, and RNA was purified using the SDS/hot-acid/phenol method. The ribosome footprint samples were resolved on a denaturing polyacrylamide gel, and a band between 20 and 45 nucleotides (nt) was excised. RNA was recovered using the ZR small-RNA PAGE recovery Kit (Zymo Research). After dephosphorylation of the 3′ ends of the recovered RNA using T4 polynucleotide kinase (T4 PNK; NEB), RNA was ligated to Linker-1 (5′-App CTGTAGGCACCATCAAT ddC-d′) using the following reaction components: 20% (wt/col) PEG, 10% DMSO, 1 × T4 Ligase reaction buffer, 20 U SUPERase·In, and 10 U T4 Ligase 2, truncated (NEB). Ligated products were resolved on a 10% TBE-Urea gel, and a band between 30 and 70 nt was excised. Ligated RNA was recovered using the ZR small-RNA PAGE Recovery kit. After phosphorylation of the 5′ ends of the 3′-ligated samples using T4 PNK, Linker-2 (5′-GAGTCTGCGTGTGATTCGGGTTAGGTGTTGGGTTGGGCCA-3′) was ligated using T4 RNA Ligase1 (NEB). Reaction mixtures were resolved on a 10% TBE-Urea gel, and a band between 90 and 120 nt was excised. Ligated RNA was recovered using the ZR small-RNA PAGE Recovery kit. cDNAs were synthesized using SuperScript III Reverse Transcriptase (Invitrogen) and Linker-1-RT (5′-ATTGATGGTGCCTACAG-3′) as a primer. RNA products were hydrolyzed by adding 1 mM NaOH to a final concentration of 0.1 mM and incubating for 15 min at 95°C. The cDNA products were resolved from the unextended primer on a 10% TBE-Urea gel, and a band between 90 and 120 nt was excised. DNA was recovered using the ZR small-RNA PAGE Recovery Kit. The resulting cDNA was PCR amplified with Q5 Hight Fidelity Polymerase (NEB) using LInker-2-partial (5′-TTAGGTGTTGGGTTGGGCCA-3′) and Linker-1 as the primers. Amplified PCR products were purified using AMPure Bead (Beackman Coulter). KAPA Hyper Prep Kits for Illumina (KK8500) were used to construct the library, and the resulting DNA libraries were sequenced using the Illumina HiSeq 1500 system. The Illumina libraries were preprocessed by clipping the adapter sequences (Linker-1 and Linker-2) using CutAdapt v.2.10 [22] for the linker sequences, and Trimmomatic v.0.39 [20] for the Illumina adapters. Then, the sequencing reads were aligned to the S1 genome sequences [7] using HASAT2 v.2.2.1 [21]. Next, the S1 gene feature file [7] was used to determine the CDS region. Sequencing data were deposited in the DDBJ database with the accession number DRA011224.

### 5.7. Definition of the ribosome density

The ribosome density (RD) was calculated as previously described [23], except that ribosome footprints between 24 and 30 nt were used for the calculation. The rationale for choosing this range was to identify as many footprints as possible to improve the statistical power and to exclude those that were suspected not to be responsible for the footprints.

### 5.8. Definition of A-site peaks

First, we determined the A-site corresponding to each read by an offset of [(15/27)×(L)] from the 5′ end of the read, where L is the length of each read. We used the normalized A-site count (ribo/mRNA) as A-site peaks.

### 5.9. Metagene analysis for ribosome stalling

To obtain the metagene profile illustrated in Figure 3B, an A-site density higher than eight was scaled by its own mean RD. Each normalized RD profile was aligned by their respective start codon and averaged across each position.

### 5.10. Definition of the stalling peptides

The stalling P-values were calculated as previously described [11]. Peptides with the P-value less than 0.0001 are termed here as stalling peptides.

### 5.11. Enrichment of over- and underrepresented peptide sequences

The over- and underrepresented peptide sequences were identified as previously described [11], except that the cutoff of the P-value was 0.0001 for the underrepresented peptides and 0.9999 for the overrepresented peptides.

## 6. Declarations

### Ethics approval and consent to participate

Not applicable.

### Consent for publication

Not applicable.

### Availability of data and materials

All data generated in this study have been deposited to the DDBJ depository (DRA011224).

## Acknowledgments

This work was funded by Grant-in-Aid for Japan Society for the Promotion of Science Research Fellow 16J02984. We thank Tomoko Yamamoto and Akiko Takaya for discussions.

## Author contributions

ST contributed to the conception and overall design of the work, all of the experiments, bioinformatics analysis, and interpretation of Ribo-Seq data and drafting and revision of the manuscript. The author read and approved the final manuscript.

## Supplementary tables

**Supplementary Table 1.** Bacterial strains.

| Strain | Relevant characteristics | Reference of source |
| --- | --- | --- |
| *Streptococcus pneumoniae* |  |  |
| S1 | TEL-resistant clinical isolate | [6] |
| Sp36 | TEL-resistant mutants of S1 isolated from 8 | [6] |

|  |  |  |
| --- | --- | --- |
|  | μg/mL TEL-containing BHI-Y agar plates |  |
| Sp52 | TEL-resistant mutants of Sp36 isolated from | This study |
|  | 128 μg/mL TEL-containing BHI-Y agar plates |  |
| Sp274 | *ΔtlrB::aad(9)* in S1 | [6] |
| Sp284 | Sp274 harboring pTKY1111 | [4] |
| Sp288 | Sp52 harboring pTKY1127 | This study |
| Sp375 | Sp52 harboring pTKY1111 | This study |
| Sp378 | Sp52 harboring pTKY1111 | This study |
| Sp379 | Sp274 harboring pTKY1127 | [4] |

**Supplementary Table 2.**
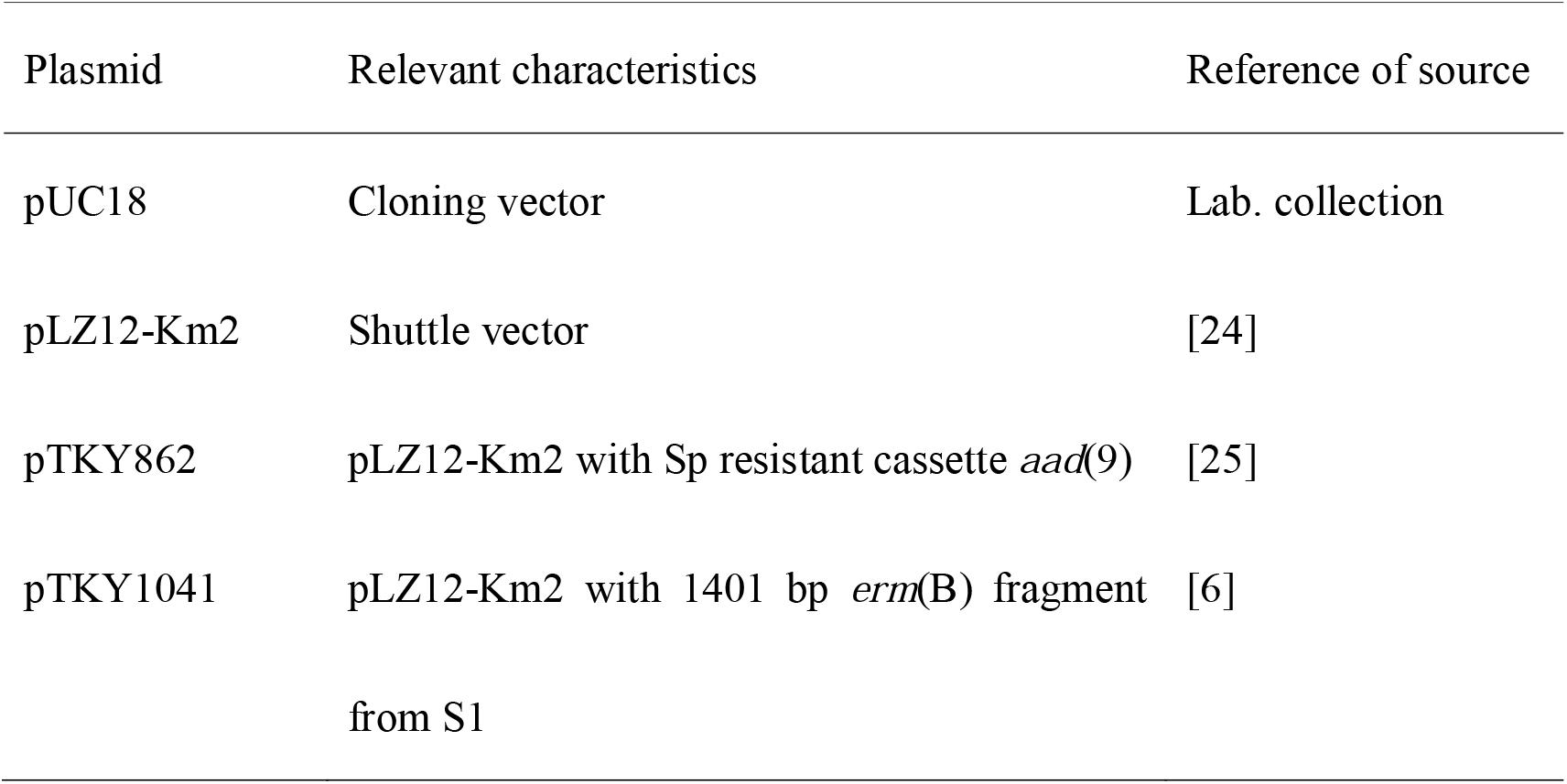
Plasmids.

| Plasmid | Relevant characteristics | Reference of source |
| --- | --- | --- |
| pUC18 | Cloning vector | Lab. collection |
| pLZ12-Km2 | Shuttle vector | [24] |
| pTKY862 | pLZ12-Km2 with Sp resistant cassette *aad(9)* | [25] |
| pTKY1041 | pLZ12-Km2 with 1401 bp *erm(B)* fragment from S1 | [6] |

|  |  |  |
| --- | --- | --- |
| pTKY1109 | pUC18 with 328 bp *tlrB* fragment | [6] |
| pTKY1110 | pUC18 with disrupted *tlrB* fragment by<br>insertion of an Sp resistant cassette *aad(9)* | [6] |
| pTKY1111 | pLZ12-Km2 with 1065 bp *tlrB* fragment from<br>S1 | [6] |
| pTKY1127 | pLZ12-Km2 with 1065 bp *tlrB* fragment from<br>Sp44 | [6] |

